# SpheroidPicker: An Automated 3D cell culture manipulator robot using deep learning

**DOI:** 10.1101/2020.11.25.397331

**Authors:** Istvan Grexa, Akos Diosdi, Andras Kriston, Nikita Moshkov, Maria Harmati, Krisztina Buzas, Vilja Pietiainen, Krisztian Koos, Peter Horvath

## Abstract

Recent statistics report that more than 3.7 million new cases of cancer occur in Europe yearly, and the disease accounts for approximately 20 % of all deaths. High-throughput screening of cancer cell cultures has dominated the search for novel, effective anticancer therapies in the past decades. Recently, ex vivo 3D cell cultures from the patient’s own cancer cells have gained importance. We recently evaluated the major advancements and needs of the 3D cell cultures screening field, and we concluded that strictly standardized sample preparation is the most desired development. Here we propose an artificial intelligence-guided low-cost 3D cell culture delivery system. It consists of a light microscope, a micromanipulator, a syringe pump, and a controller computer. The system performs morphology-based feature analysis on spheroids and transfers the most appropriate ones between various sample holders. It can select the samples from standard sample holders, including Petri dishes and microwell plates, and then transfer them to a variety of holders up to 384 well plates. The device performs reliable semi- and fully automated spheroid transfer. This results in highly controlled experimental conditions and eliminates non-trivial side effects of sample variability that is a key aspect towards next-generation precision medicine.

## Introduction

In drug discovery studies, cell-based assays have been widely adopted as a model system. In order to understand the activity of a compound, two-dimensional (2D) model systems are still in use (Horvath et al., 2016). However, it has been shown that the microenvironment of the in vivo tumor is different from the 2D monolayer cell cultures (Hoarau-Véchot et al., 2018). It has recently been described that three-dimensional (3D) cell cultures are more relevant model systems for drug testing (Carragher et al., 2018) because they enable the examination of drug penetration and tumor development (Sawant-Basak & Scott Obach, 2018; Szade et al., 2016). One of the most common 3D cell cultures is the spheroid models where the cells form a sphere-like structure (Cesarz & Tamama, 2016). Although spheroid models can reflect in vivo conditions better, it is still a challenge to use these models in a high-throughput system. The lack of a unified protocol for creating spheroids results in varying shapes with huge morphological heterogeneity (Cisneros Castillo et al., 2016) that is not ideal for clinical studies. In most cases pre-selecting spheroids by their morphology is a very time-consuming and inconsistent process because it requires human assistance. Moreover, the expert selects the spheroids by the naked eye or in the case of smaller samples (under 500 µm in diameter) under a simple microscope.

Imaging of 3D cell cultures is limited because of the light scattering due to their compact structure (Hoarau-Véchot et al., 2018). The screening plate can greatly affect the image quality. For example, U-bottom shaped plates are suitable for growing spheroids because they have more beneficial, spherical morphologies, but negatively affect imaging quality. Therefore, it might be required to transfer these spheroids to a flat surface, e.g. a flat bottom plate or a Petri dish.

Spheroids derived from cell lines and patients will result in different shapes and sizes that can make the selection difficult. Therefore, detection and segmentation must be precise to achieve highly accurate features. Previous selection techniques are based on subjective decisions of experts, who specified the optimal conditions, such as area or perimeter (Bresciani et al., 2019). Unfortunately, the imaging conditions can strongly vary, such as light, illumination, density of the medium, or the shape of the plates that can affect the expert’s decision. Spheroids should be detected and segmented on label-free brightfield microscopy images. Methods that handle such images are based on classical analysis methods like thresholding or watershed (Bresciani et al., 2019; Carpenter et al., 2006). Most recently, deep learning object detection and segmentation methods have become popular since they provide more accurate results, and do not require fine-tuning even if imaging conditions change (Hollandi et al., 2020).

Existing commercial devices are able to automate various parts of cell culture handling, for example, automatic cell generation with Sphericalplate 5D (Doulgkeroglou et al., 2020). These regulate the spheroid size, which is just one step for strandization. Most of the systems support liquid treatment but not selective transferring. Devices created for plate to plate transferring (Yamaha Cell Handler) do not use advanced cell detection methods and always expect one cluster or spheroid per plate. Methods used for detection are threshold-based methods, which can lead to inappropriate feature calculation, e.g. regarding the size of the cell culture. These methods usually utilize machine learning only for cell classification, but not for segmentation.

In this work, we propose a new solution to work with spheroids. We designed and built a robotized microscope that can support or fully replace experts by automatically selecting and transferring 3D cell cultures for further experiments (Fig. 1). We developed a fast and accurate deep learning-based framework to detect and segment spheroids and integrated it into the machine’s controller software. We have created a unique image database of annotated spheroids and used it for training deep learning models. We show that our models can detect and segment spheroids with high reliability. Using such accurate segmentation, the algorithm is able to extract features robustly and - based on the user’s criteria - decide which spheroids to manipulate. This system was developed to automate one of the most time-consuming processes during sample preparation, the pre-selection phase, where accuracy is crucial.

**Figure 1.**
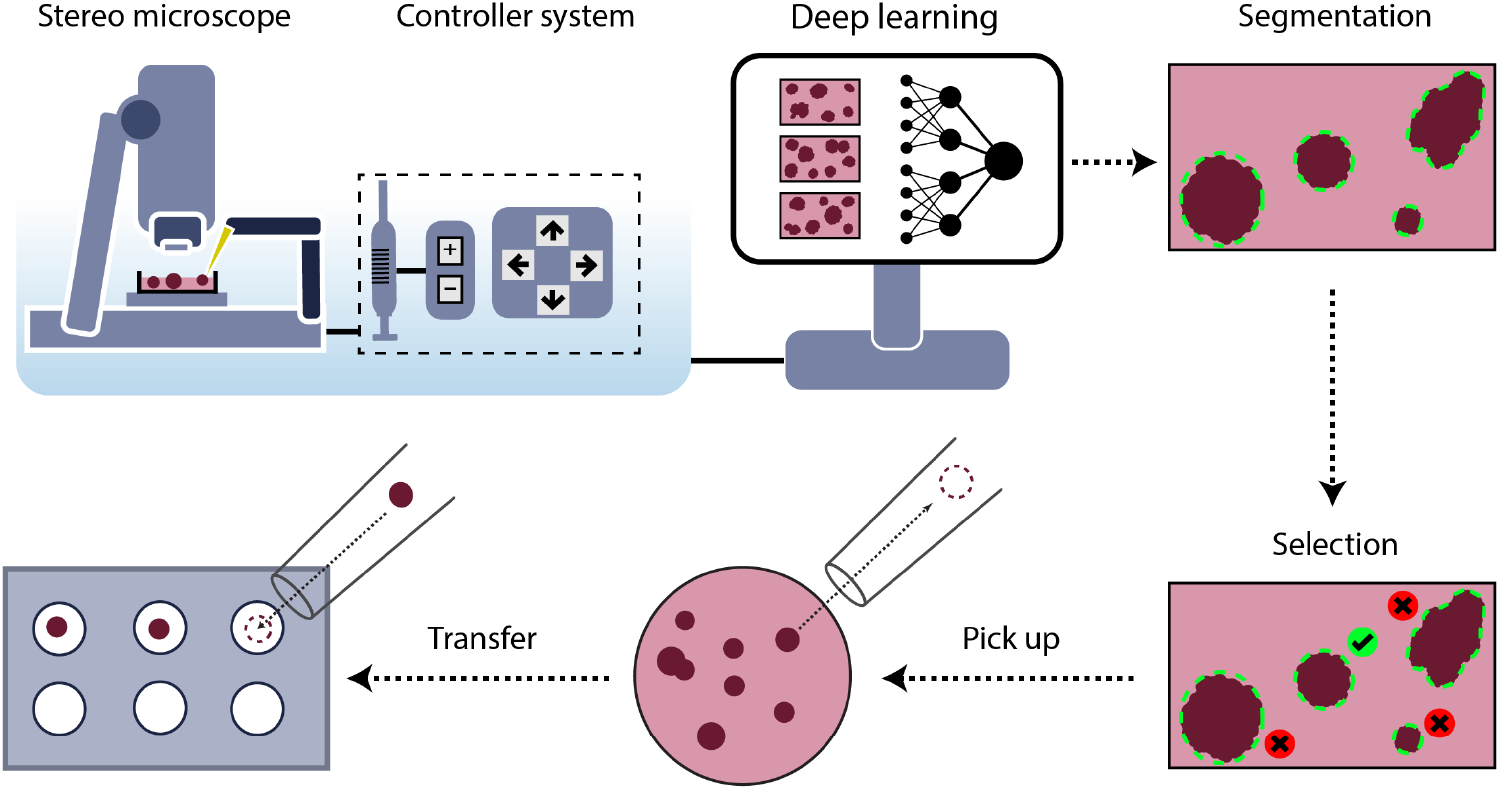
Schematic representation of the SpheroidPicker. The system includes a stereo microscope, syringe, stage, and manipulator controller. The automatic screening function results in brightfield images of the spheroids. The segmentation and feature extraction steps are based on a deep learning model. After the selection of the spheroids, the spheroid picker automatically transfers the spheroids into the target plate.

## Materials and methods

### Components of the system

The system is built on a stereomicroscope (Leica S9i, Germany) with a large working distance to fit the pipette between the sample and the objective. This microscope has an integrated 10MP CMOS camera directly connected to the controller PC via USB port. The motorized stage (Marzhauser, Germany) has a 100 x 150 mm movement area. We designed a 3D printed plate holder that is compatible with the most common microwell plates, including 24-, 96-, or 384-well plates (Fig. 5 (c-d)). A two-axis micromanipulator is installed next to the microscope and can be controlled at a micrometer precision. A custom-designed stepper motor-driven syringe pump ensures high precision (3 µl) fluid movement. The whole system is small, thus mobile, and can be installed under a sterile hood.

The software performs four major operational steps 1) entire hardware control, 2) automated imaging, 3) spheroid detection using a trained Mask R-CNN (He et al., 2017) deep learning framework (described later), and 4) an easy to use graphical user interface to define custom spheroid selection and transfer criteria.

### Custom micromanipulator and syringe pump

The custom micromanipulator is designed and built based on the openbuilds (OpenBuilds Team) linear actuators. The mainframe is built from 2080 (C-beam) aluminum extrusion. An M8 threaded rod with 2 mm pitch moves a four-wheel carriage. A Nema 17 two-phase bipolar stepper motor is used to directly rotate the threaded rod which provides enough torque and holding force. Each actuator has two endstops that are simple switches, and when they are hit with the carriage they help to avoid collision. These also guarantee the precise initialization at startup by setting these positions as the origin. Two of these actuators are assembled at 90 degrees, one of them provides axial positioning, the other sets the height of the pipette. The pipette is a glass capillary rod that is inserted into a capillary holder and attached to a 3D printed part.

The custom syringe pump is a simplified linear actuator that is moved by a stepper motor. It consists of 3D printed parts. Sterile or disposable syringes can be used to ensure the sterility of the system. The syringe is connected to the capillary holder with a silicone tube. This tube has to be filled with liquid (without air) beforehand to ensure precise flow. The parameters of the capillary are 1 mm OD, 0.6 mm ID, and 40 mm in length. The hardware is controlled with an Arduino Mega 2560 microcontroller combined with “ramps 1.4” shield. This shield provides routes and pinouts from the Arduino which makes it easy to connect stepper motors, drivers, and high power.

The Marlin open-source CNC controller software is used as firmware. It operates with G-code commands that are sent via serial port. The framework has many additional features besides simple hardware driving, such as speed, acceleration, and jerk control.

### 3D printing

Parts were 3D printed with an unmodified Original Prusa i3 MK3s, which is an FDM cartesian type 3D printer. It melts the spooled thermoplastic with a hot end on the printhead and is deposited on growing work. The head is moved with a small computer control. The plate holders were created with PLA, printed with 60 °C bed temperature and 210 °C hotend temperature. The parts of the manipulator were printed with PETG 90 °C bed temperature and 245 °C hotend temperature. The models were designed in Autodesk Inventor and sliced with PrusaSlicer. The printer has the default installed with a brass 0.4 mm hole diameter nozzle, and we used 0.15 mm layer height.

### Software

The main controller software is written in C++. It has an easy to use GUI implemented in QT framework (Fig. 2). The goal of the software is to operate all the hardware components through serial connections that are the digital camera, stage, and micro-pipetting system.

**Figure 2.**
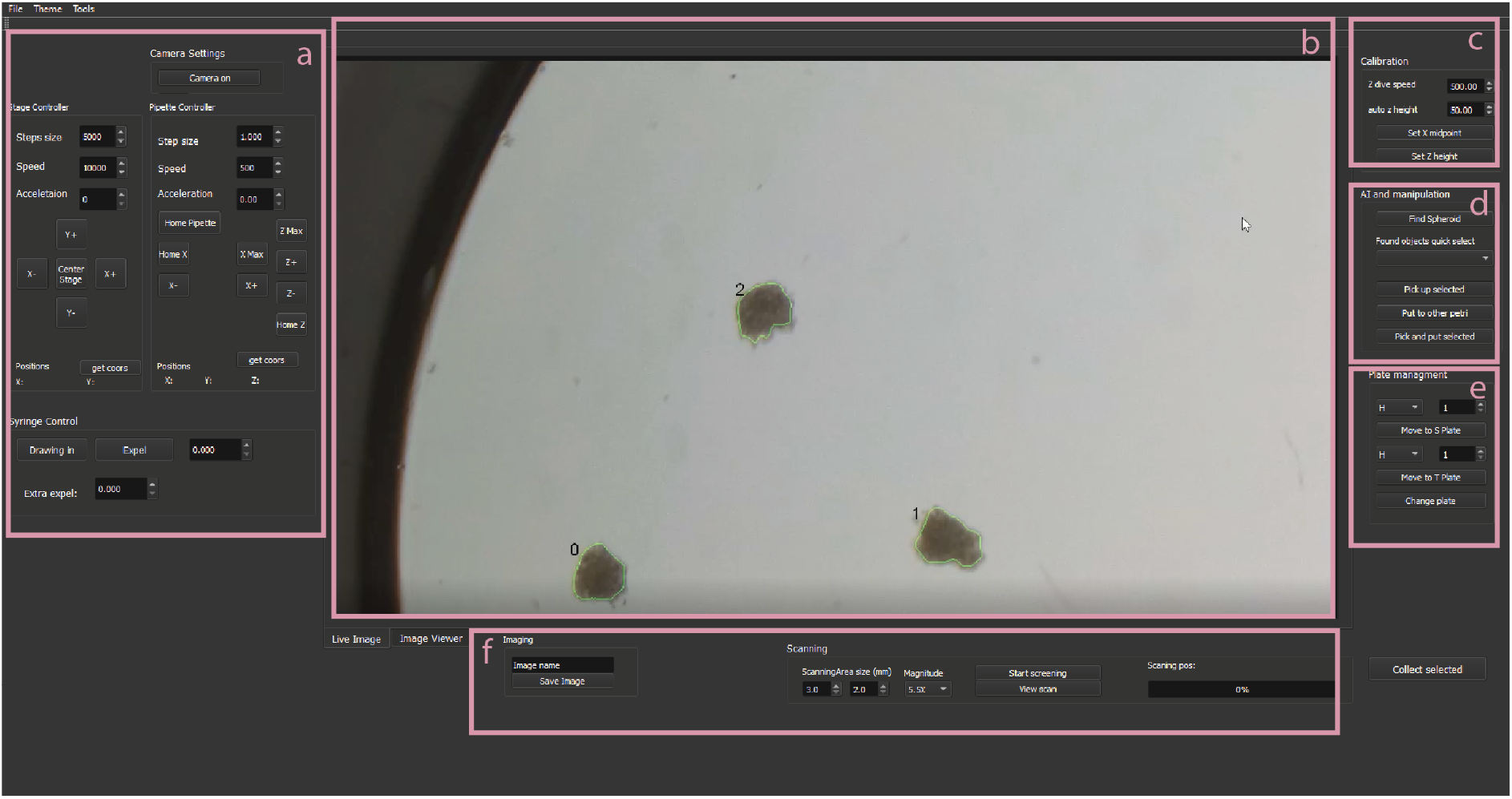
The main window of the software. The user can access the hardware control, imaging, and spheroid detection system. It can be used for spheroid manipulation, plate management, and scanning. **a:** Hardware control tools, e.g. pipette or stage movement, or syringe dosage. **b:** live or segmented image. c: Picker calibration tools. **d:** Semi-and fully automatic spheroid manipulation tools. **e:** Buttons for a quick jump to wells. **f:** scanning tools.

The system has to be calibrated before the first use to be able to move the pipette based on image coordinates. The process can be initiated from the software and consists of the following steps. First, the user puts a well plate in the holder and safely moves the pipette under the objective. Then the pipette tip should be moved as close to the surface of a well as possible to determine the lowest height without crashing into the plate. Finally, the tip should be moved to the center of the image and register the coordinates in the software. When calibrated, the system works with every supported plate. The process should take less than 5 minutes.

The software includes a Mask R-CNN deep learning model trained to segment spheroids. Input images are downscaled to 1024 × 1024 pixels by keeping the aspect ratio and zero-padding. The model is developed with TensorFlow/Keras libraries, the images are handled with OpenCV libraries. After segmentation, the main features of the objects are calculated, including area, perimeter, circularity, mass center, and stage coordinates. With a mid-range GPU (NVIDIA 1050 Ti 4Gb GDDR5, 768 CUDA cores @1430 MHz) the inference takes about 1 second per image.

The software can scan the supported plates and select target spheroids that satisfy the requested properties. When the scanning finishes, the transfer protocol can be initiated. The stage centers on one of the selected objects in the camera image, the X-axis (effector) moves the pipette above it, and Z-axis lowers close to the well surface. A negative flow is applied with a syringe pump that soaks up about 3-4 μl fluid with the target spheroid. Afterward, the pipette is lifted and the stage moves the target plate and well under the objective. Then the pipette is lowered and the spheroid is pushed out to finish the transfer. Finally, the system either starts over the process with the next spheroid or pulls out the pipette of the field of view to finish the session.

### Cell cultures

T47-D human breast cancer cell line was cultured in RPMI-1640 (Lonza, Switzerland) supplemented by 10% FBS (EuroClone, Italy) and 0.2 unit/ml insulin (Gibco, Thermo Fisher Scientific, USA). Huh-7D12 human hepatocellular carcinoma cells were cultured in DMEM (Lonza, Switzerland) supplemented by 10% FBS and 1% L-glutamine (Lonza, Switzerland). 5-8F human nasopharyngeal carcinoma cell line was cultured in DMEM-F12 (Lonza, Switzerland) supplemented by 10% FBS. A549 human lung carcinoma cells were cultured in RPMI-1640 supplemented by 10% FBS. All media contained 1% Penicillin-Streptomycin-Amphotericin B mixture (Lonza, Switzerland) as well. Cell cultures were maintained at 37°C and 5% CO_2_ in a humidified incubator.

### Spheroid generation and fixation

Multicellular spheroids were created by SphericalPlate 5D (Kugelmeiers, Switzerland) based on the manufacturer’s instructions. To seed 750 cells for each microwell/spheroid, we applied 5.625 ×10^5^ cells in 2 ml medium per well. The spheroidization time was 7 days for the T47-D, 5-8F, and A549 cell lines, and 4 days for the Huh-7D12 cells to reach similar spheroid diameters ranging from 200 to 250 µm. Culture media were changed every other day during the incubation time. After the spheroids developed, they were washed twice with DPBS and fixed with 4% paraformaldehyde for 1 h at room temperature. Finally, the spheroids were washed twice with DPBS and stored at 4°C in DPBS.

### Data Generation

We have acquired 981 brightfield images of spheroids in very different illumination conditions. These images (1920 × 1080, 24 bit RGB) were annotated by expert biologists using ImageJ (Collins, 2007) to generate mask images as ground truth data for the training of the model. Finally, we had 1871 annotated objects. We divided these images into 3 sets: 70% for training, 15% for validation, and the rest 15% is used for testing.

### Segmentation methods

Precise image segmentation is the foremost important step of our spheroid selection pipeline. We measured the precision and sensitivity of four different widely used segmentation methods. Classical Otsu thresholding and watershed-based segmentation were benchmarked using the CellProfiler framework (CP) (Carpenter et al., 2006). Furthermore, U-Net (Ronneberger et al., 2015) and Mask R-CNN (He et al., 2017) deep learning techniques were adapted to the problem. CellProfiler (Carpenter et al., 2006) is a widely-used bioimage analysis software including methods to segment, measure, and analyze cellular compartments. We have created 2 pipelines: for primary object detection one uses the Otsu thresholding method and the other uses the watershed method. Both pipelines have additional filters to remove the objects which have a too small or too large area or have an irregular shape. Important to note that, the CP parameters were set to get the best results on all the test images, not for images individually.

The U-Net neural network is designed for biomedical image analysis (Ronneberger et al., 2015) and in total has 23 convolutional layers. It is designed for semantic segmentation which does not handle overlapping objects. However, since spheroids usually do not touch or overlap, we consider the network appropriate for this problem. We trained U-Net for 100 epochs with learning rate (LR) = 3×10^−5^ using Adam optimizer. The model was trained on a PC with Intel Xeon CPU E5-2620 v4 @ 2.10GHz CPU, Nvidia Titan Xp 12GB VRAM GDDR5x (founders edition, reference card), 3840 CUDA cores @1544Mhz with Pascal architecture, 32 GB DDR4 of RAM.

Mask R-CNN is a general framework for the instance segmentation task (He et al., 2016). We trained Mask R-CNN with ResNet101 backbone initialized with COCO weights. The training consisted of 4 stages and 100 epochs in total, where different layers were updated and the learning rate (LR) was changed. In the first 4 epochs, we updated every layer in the ResNet with LR = 5×10 ^−4^. In epochs 5-7 only 5th and above layers were updated with LR = 3×10 ^−5^, and finally only the network head layers with LR = 1×10 ^−7^. LR values were empirically selected based on the loss function values. The training, performed on the same system as U-Net.

## Results

Here, we describe how we developed and built hardware and software solutions (Materials and methods) to automate the spheroid handling process between various well plates (Fig. 3). The main parts of the hardware are a stereo microscope, a micromanipulator, a syringe pump, and a computer (Fig. 1).

**Figure 3.**
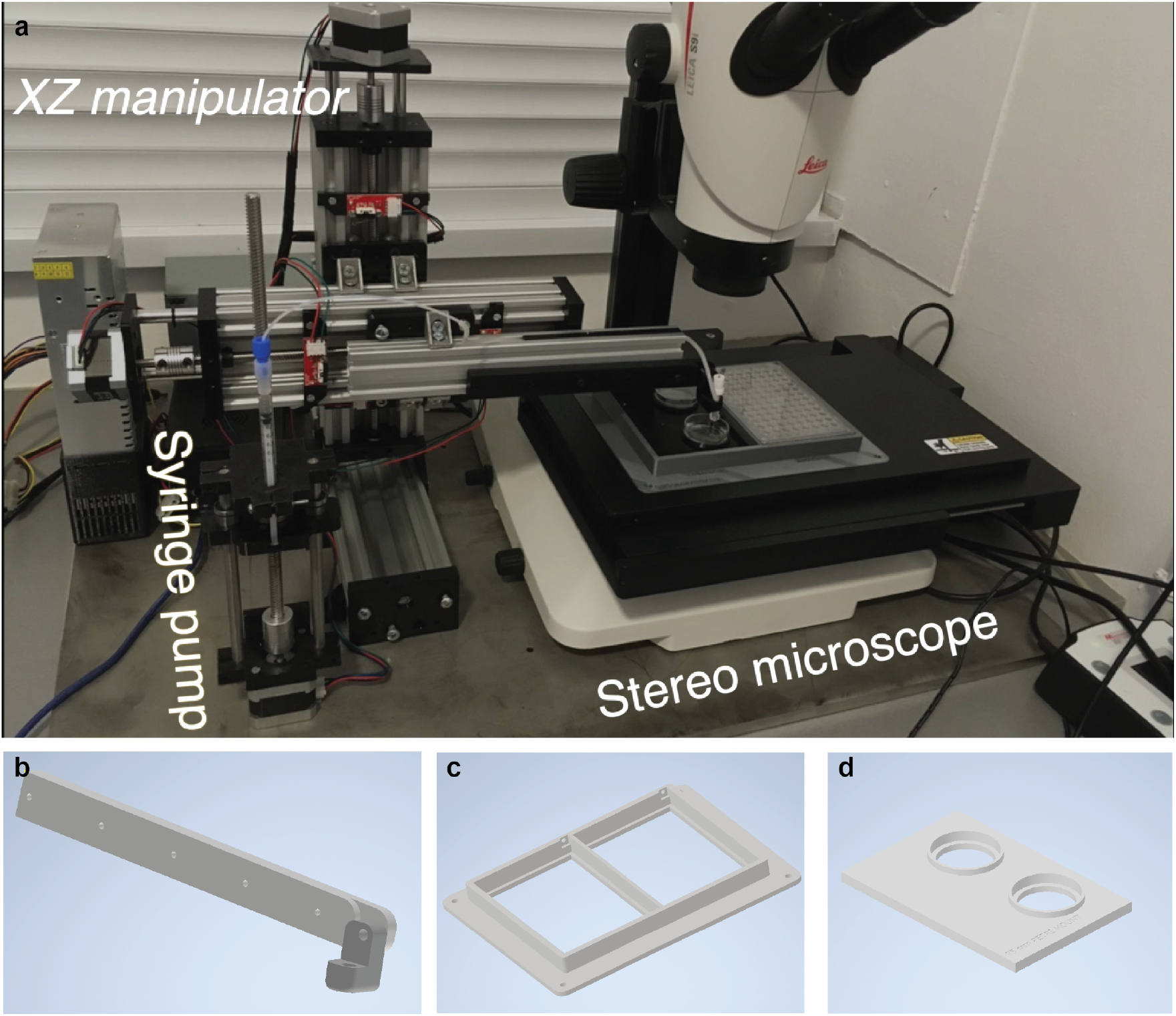
**a:** SpheroidPicker prototype. The main components are shown, which are the manipulator, the syringe pump, and a stereo microscope. **b:** The 3D printed element that fixes the capillary holder to the linear actuators. **c:** 3D model of the custom-made plate-mounting system that is compatible with the most common 24-, 96-, 384-well plates. The source and target plate can be placed next to each other. **d:** model of a petri dish holder which can be inserted into the 96 well plate holder.

The large field of view of the microscope allows for the efficient and fast screening of the samples. Various well plates and Petri dishes are supported so that wells or regions can be selected or excluded from analysis. The movement area of the motorized stage is large enough to place the source and target plate next to each other, and switch between them easily. The micromanipulator moves a glass capillary rod that is used for transferring the spheroids. Its axes can be configured to move it based on image coordinates. The capillary is further attached to an automated water-based syringe system. Except for the microscope, the stage, and the camera, all the hardware elements are custom built. The components are controlled from a single, custom software running on a computer. The device creates a standardized protocol for spheroid handling which is absolutely necessary for drug screening experiments.

A key feature of the system is the automatic spheroid selection capability. The operator can specify morphological properties that are required for the selected objects, such as a size range or circularity value. For the calculation of these values, a reliable image segmentation method is needed. We compared four segmentation methods which are Otsu thresholding, watershed algorithm, U-Net, and Mask R-CNN. In general classical image analysis methods could be used for the segmentation task, however, they are rather sensitive to changes in imaging conditions such as light, medium density, object transparency, shape diversity, and touching objects. Another disadvantage of these methods is that the user has to reconfigure parameters at every experiment, which can lead to loss of morphological features or getting false results. On the other hand, deep learning methods have gained attention during the last few years due to their learning capabilities and robustness. We show that our deep learning models can reliably predict spheroids without reparameterization in every condition even if the image is out of focus.

We trained a Mask R-CNN and U-net deep learning framework to segment the spheroids. The learning curves are shown in Fig. 4. We used the multitask loss function defined in the Mask R-CNN architecture which combines the classification loss, localization, and segmentation. For U-net we used binary cross-entropy loss functions.

**Figure 4.**
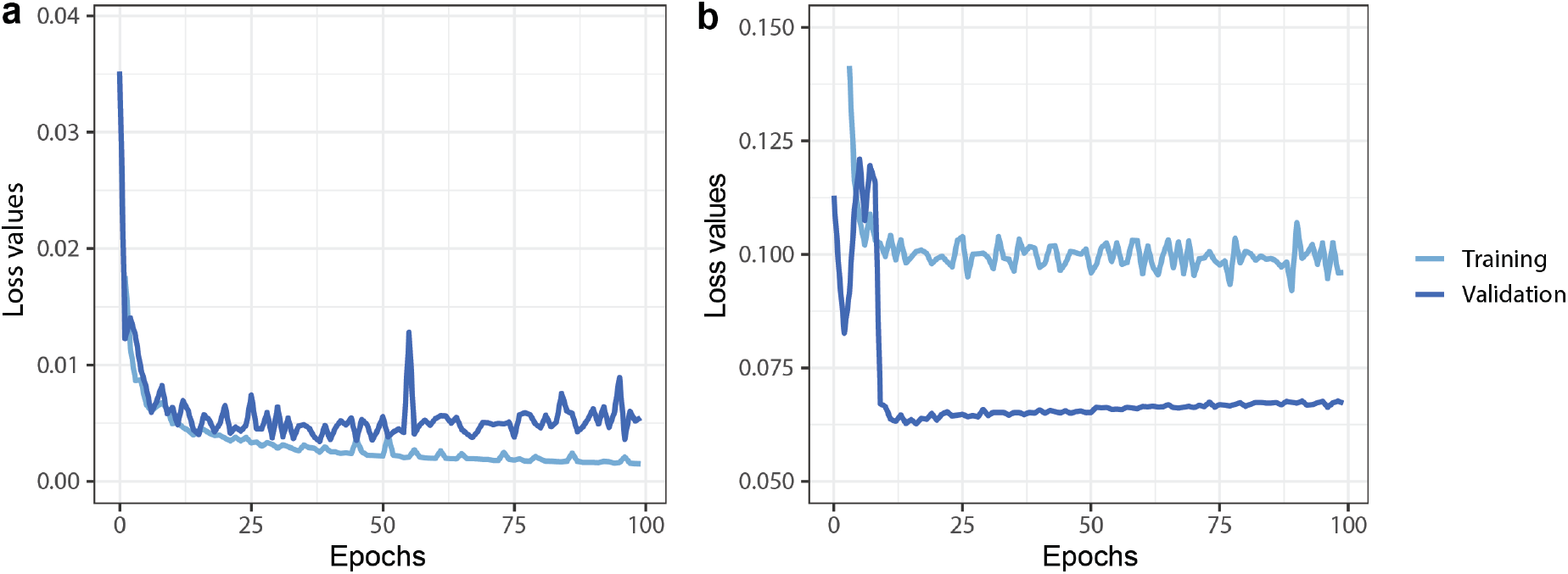
Deep learning training and validation loss values through 100 epoch training. **a**: U-net, **b**: Mask R-CNN. Both Neural networks are convergent, with small training and validation loss values.

The model’s performance was measured on a test set that consisted of 153 images. We selected in both Mask R-CNN and U-net the model with the lowest validation values. First, we analyzed the quality of the 4 segmentation methods. The results of the classical methods and the deep learning model were compared to the expert annotations (Fig. 5). We concluded that one classical segmentation pipeline cannot give reliable results in different experimental conditions. Although some spheroids are recognized remarkably well, in most cases it fails to find precisely spheroids (Fig 5b). Deep learning methods give very small false results, and contours almost pixel-wise precise. We can observe some cases where the U-net is not able to completely differentiate two very close spheroids (Fig. 5d). However, the contours produced by U-net are usually closer to the annotated, than Mask R-CNN (Fig. 5a) which is also confirmed by precision comparison (Fig. 6).

**Figure 5.**
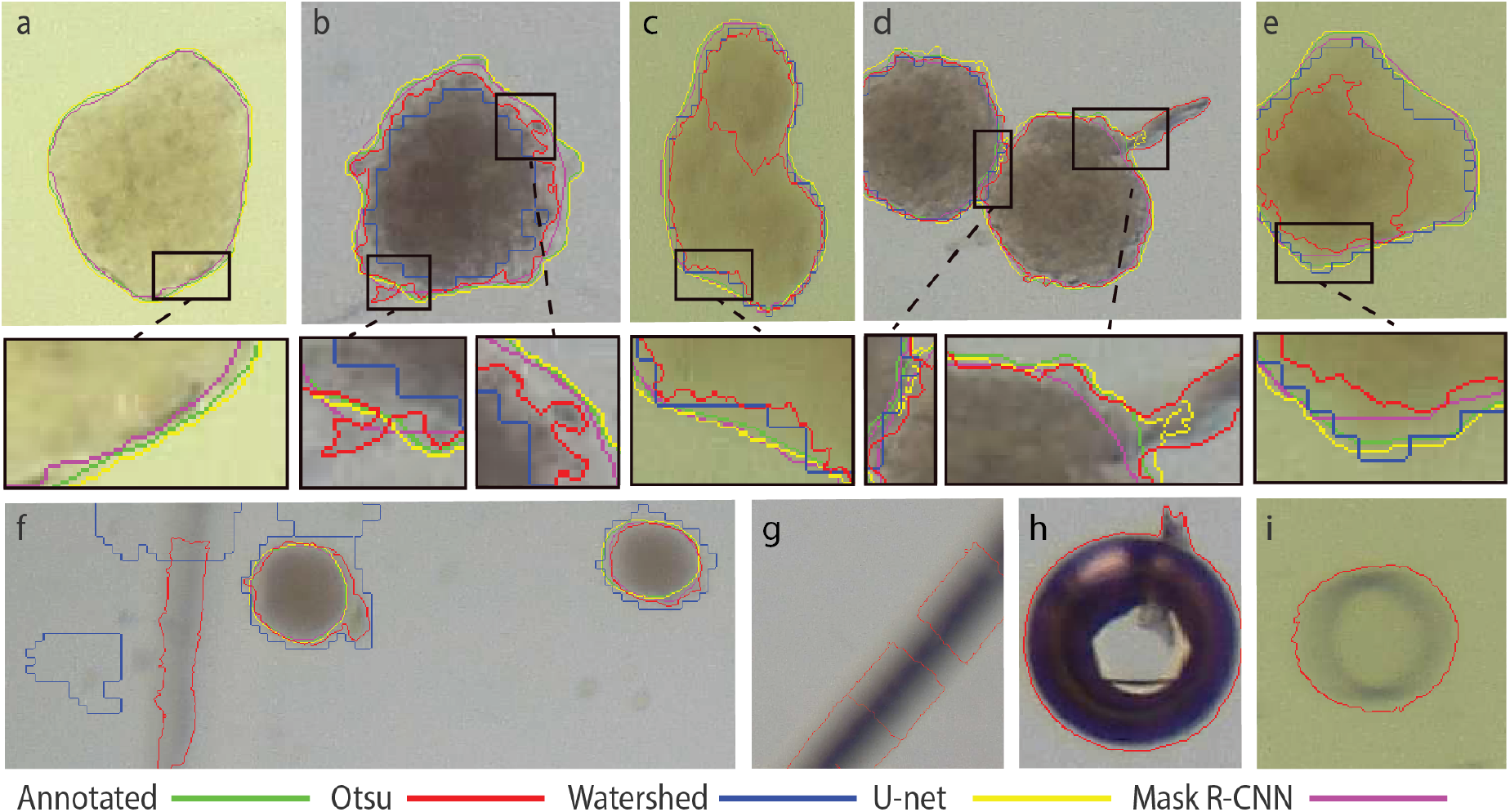
Brightfield images of spheroids with various shapes and sizes. Images contain results of 4 different segmentation methods and the annotated ground truth with color codes: green - ground truth; red: Otsu threshold; blue: watershed; yellow: U-net; magenta - Mask-R-CNN. **a-e**: Selected results and enlarged regions. In **a** only deep learning methods could find the spheroid. In **d** we can observe that the U-net is not able to completely differentiate two very close spheroids. In **c**, thresholding identifies one spheroid as two because of the intensity changes. **f-i**: artifacts made by thresholding and watershed. These are usually out of focus spheroids, borders of the plates, plastic errors in plates, or air bubbles.

**Figure 6.**
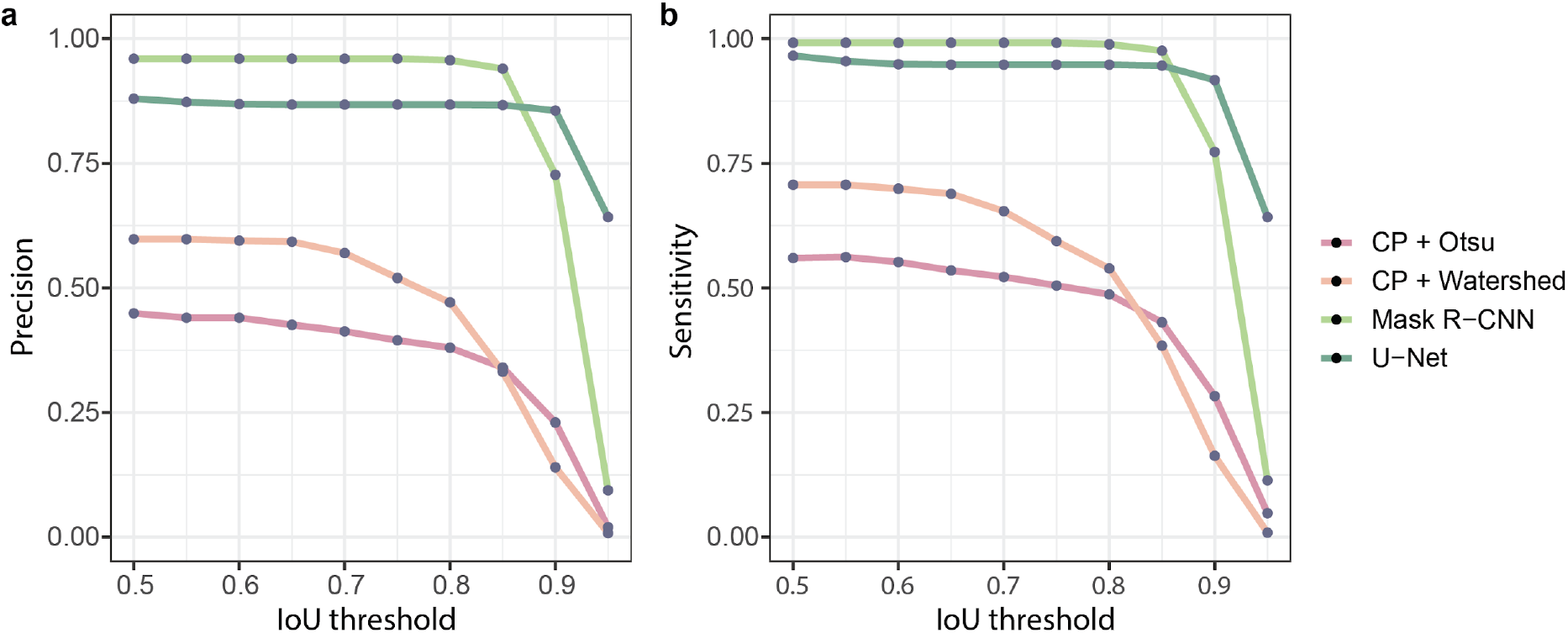
Comparison of the segmentation methods. **a:** average precision, **b:** sensitivity scores, at the different intersections over union thresholds.

We compared the described segmentation methods by measuring the average precision (P) and sensitivity (S) scores (Eq. 1) on different IoU threshold values (Fig. 6). Classical segmentation methods give us worse AP and S values, because of the changing light conditions and heterogeneous shapes of spheroids. Regarding deep learning methods, Mask R-CNN with IoU value between 0.5 and 0.85 performs better in terms of average precision and sensitivity scores than U-Net. This means that Mask R-CNN detects the objects more reliably, but segments them less accurately. We can see that U-Net results in better AP and S on higher 0.9-0.95 IoU values, which means it is better for pixelwise object segmentation. As we use those results for object manipulation too, accurate object detection is more important, than slightly more accurate pixel wise segmentation.

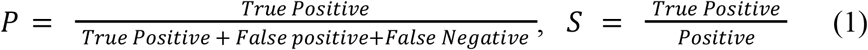

### Quality assessment

We identified and measured different types of errors the system makes. The first type is when the target object is not sucked properly into the capillary or it falls out before getting to the target well (pick up error). The second type is when the spheroid is present in the capillary but the system is unable to put it in the target well, usually due to adherence (expel error). In the third error type the system picks up two or more spheroids, which happens when these are very close to each other and one of them is the target (double pick).

The spheroid transferring capabilities of the system were tested by moving them one by one from a source 96 well plate to a target plate of the same type (Supplementary Video 1). The selection criteria was an area range between 21,000 and 29,000 µm^2^ and a minimum of 0.815 circularity. A minimum circularity criterium ensures that the selected object has a rounded shape. First, we performed this experiment such that the outcome was evaluated right after every transfer and possible issues were fixed (e.g. removed spheroids that stuck in the capillary) to be able to continue. From 28 attempts, 26 spheroids were successfully picked up, 25 were transferred properly, and always one object was picked up, which leads to a success rate of 89% (Fig. 7c). Next, we let the system fully automatically select and transfer spheroids between the plates. The Picker scanned a user-defined area of the sample and analyzed it. Out of 30 attempts, 24 were successful which leads to an 80% success rate.

**Figure 7.**
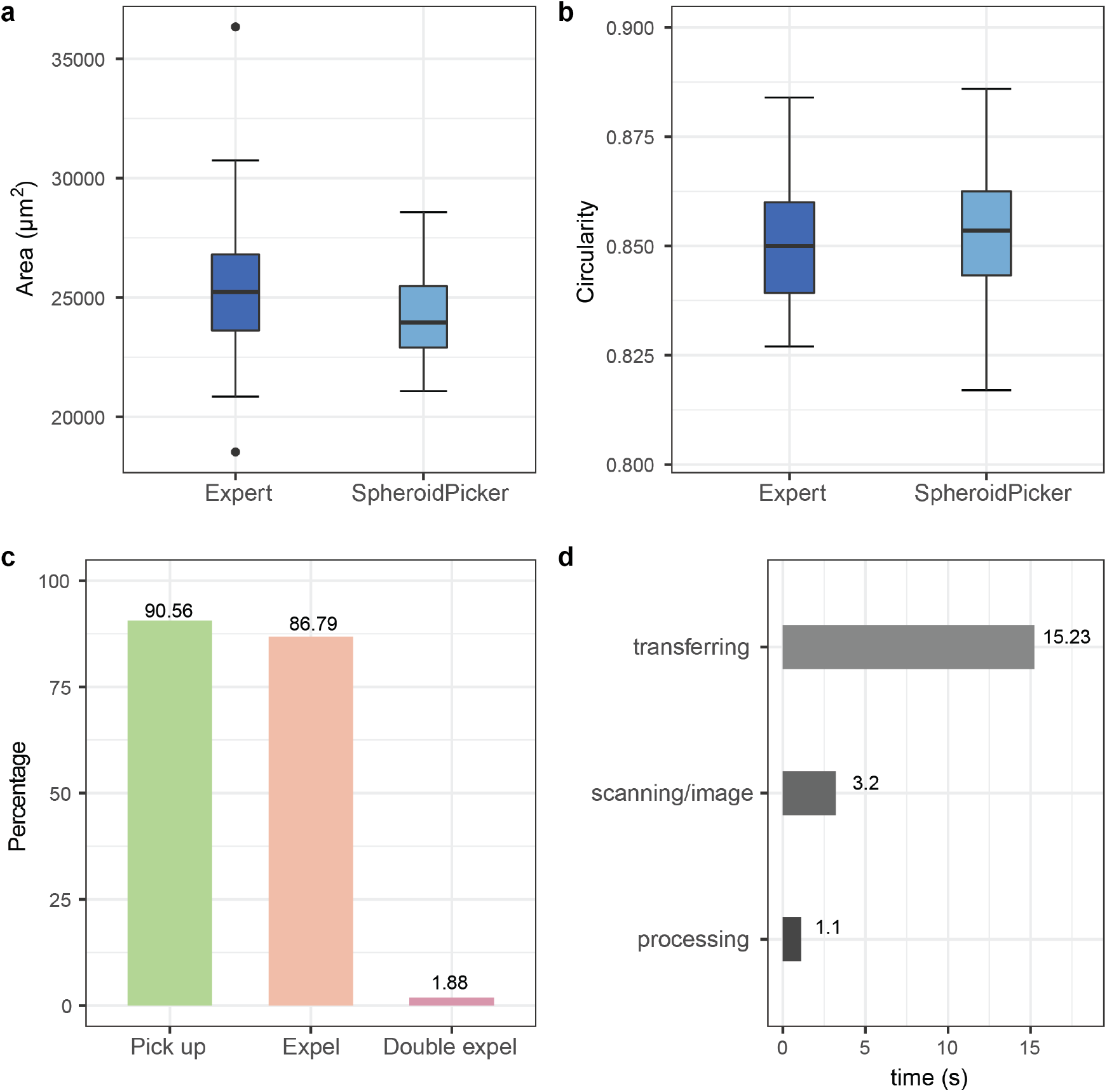
Benchmarks of the SpheroidPicker. **a-b:** Precision comparison between the expert and the SpheroidPicker. The expert selected spheroids with similar size range under the stereomicroscope. For the SpheroidPicker, 21,000-29,000 µm^2^ area range and higher than 0.815 circularity were used as selection criteria. **c:** Statistics of SpheroidPicker’s reliability. **d:** Run times of the SpheroidPicker’s tasks.

We compared the selection capability of the Spheroid Pickers to the manual process of our expert with the following experiment. First, we asked the expert to manually select preferably circular spheroids with an area in the range of 21,000-29,000 µm^2^ (preferably 25,000 µm^2^, or with 178.41 µm diameter) under the microscope with a regular laboratory pipette, and transfer them separately to the wells of a 96-well plate. Then we set the same requirements in the software and transferred spheroids using it in the order it offered them. Both the expert and device transferred 42-42 spheroids. Afterward, we used our system to determine the size and circularity properties of the manually transferred objects. The average area was 25,347.2±3,226.3 µm^2^ in the manual case and 24,222.6±1,957.7 µm^2^ in the automatic, while the circularity value was 0.8504±0.0147 and 0.8527±0.0150, respectively (Fig. 7a-b). We also measured the average time of different processes of the picker with the software: transferring, scanning and image processing (Fig. 7d).

## Conclusion

We have designed and built an automated microscopy system that is capable of selecting and transferring spheroids. The selection and the feature extraction of the spheroids are based on deep learning methods. We created to our knowledge the first annotated image database to train models for spheroid segmentation. The image processing methods used for the detection and selection are reliable, and we have shown that it can outperform human selection skills. The machine can perform semi- or fully-automatic transferring of spheroids to any predefined well plate. The greatest advantage is that it helps in selecting similar spheroids to standardize the pre-selection phase and thus make it easier and more robust. Comparing the expert and the SpheroidPicker selection capability shows that the manual approach has 1.648 higher standard deviation in terms of area, and thus our system is more reliable when specific features are required. However, the software can be improved by parallelizing scanning and image analysis tasks to speed up the process. Pick up failures and double pickups could be avoided or corrected with object detection methods, thus getting better reliability.

In conclusion, we have shown that building a microscopy system that combines AI and robotic techniques could improve efficiency in laboratory work with spheroids. To our knowledge this is the first deep learning-based manipulator robot that is designed for spheroid transfer. Compared to other devices this machine costs significantly less. The spheroid picker uses a smaller area since the main components are a stereo microscope with a stage and the custom manipulator. It uses advanced deep learning-based methods for detection segmentation.

## Acknowledgments

This project has received funding from the ATTRACT project funded by the EC under Grant Agreement 777222. We acknowledge support from the LENDULET-BIOMAG Grant (2018-342), from the European Regional Development Funds (GINOP-2.3.2-15-2016-00006, GINOP-2.3.2-15-2016-00026, GINOP-2.3.2-15-2016-00037), from H2020-discovAIR (874656), and from Chan Zuckerberg Initiative, Seed Networks for the HCA-DVP. We acknowledge support from the EFOP 3.6.3-VEKOP-16-2017-00009 grant. The authors would like to thank Ji Ming Wang (National Cancer Institute-Frederick, Frederick, MD, USA) for kindly providing us with the 5-8F cell line. We thank the NVidia GPU Grant program for providing a Titan Xp.

## Author contributions

IG built the prototype and software, AD made spheroid annotation, and experiments with the SpheroidPicker, KB and MH made the spheroids, NM and AK carried out the deep learning framework, PH, KK and VP conceived the project. All authors participated in writing the manuscript.

## Competing interests

The authors declare no competing interests.

## Data and materials availability

Annotated data is available on request.

## Software is available

Software data is available on request.

## Notes

### Competing Interest Statement

The authors have declared no competing interest.

